# Knowledge Graph and Machine Learning Help the Research of Drugs Aimed at Neurological Diseases

**DOI:** 10.1101/2024.11.29.626076

**Authors:** Luca Menestrina, Maurizio Recanatini

**Author notes:** Corresponding author. *E-mail address:* (L. Menestrina). Chemotargets, S.L. Barcelona Science Park (PCB) Baldiri Reixac 4, Barcelona 08028, Catalonia, Spain.

## Abstract

In this study, we present PATHOS (PATHologies of HOmo Sapiens), a semantically rich knowledge graph constructed by integrating diverse datasets spanning multiple biomedical entity types. PATHOS provides a comprehensive resource for representing and exploring the intricate relationships underlying human diseases. To leverage this resource, we developed LOGOS (Learning Optimized Graph-based representations of Object Semantics), a graph embedding model capable of generating predictions relevant to drug research. The PATHOS-LOGOS framework was validated through three neurological case studies: drug repurposing for Alzheimer’s disease, phenotype selection for Huntington’s disease, and protein target identification in multiple sclerosis. The results demonstrate the potential of this approach to advance therapeutic insights and inform biomedical research.

## 1 Introduction

Network theory represents a potent analytical tool for exploring complex systems, particularly in the context of living organisms. Embracing the principles of network science, this approach captures the holistic behavior of a system and highlights emergent properties that result from intricate interactions among its components.[1] This methodology is useful in understanding the pathophysiological basis of diseases, conceptualized as emergent properties arising from complex interplays within living systems.[2] Recently, network models have played a pivotal role in characterizing drug-disease relationships and shedding light on various aspects of drug research.[2] These models depict nodes representing entities connected either physically or conceptually, forming a dynamic framework to untangle the intricacies of these complex systems.[3]

In 1957, C. H. Waddington diverged from the prevailing Mendelian “one gene - one phenotype” model, pioneering a new perspective that highlighted the influence of gene networks on cellular states and develop-mental outcomes.[4] He proposed that phenotypes arise from stable conditions (states) of gene networks and the transitions between them.[4, 5] This concept lays the foundation for network biology, a discipline dedicated to unraveling the complex interactions among biomolecules constituting vital functional units driving physiological functions at the cellular, tissue, and organ levels.[6, 7]

A parallel transformation can be observed in the traditional target-centered approach to treating diseases, which traces its origins to the pioneering work of Paul Ehrlich and his renowned statement *“Corpora non agunt nisi fixata”*.[8] Despite being inherently reductionist, the “one disease-one target-one drug” model drove scientists to focus on molecules and their interactions, and served as the foundation for much of the research that led to the development of many of the medicines used nowadays.[2] Nevertheless, it has become increasingly evident that numerous effective drugs exert their effects on multiple targets rather than a singular one, a phenomenon called polypharmacology.[9] Indeed, the integration of systems biology and the expansion of the “omics” technologies has prompted a shift in the drug discovery paradigm towards what is now known as network pharmacology.[2, 9]

This strategy finds its place within the framework of network medicine, an established discipline rooted in the application of principles of network theory and network biology to disease mechanisms and pharmacotherapy [10, 11]. The key concept of network medicine is that the comprehension of drug actions and the design of innovative pharmacological treatments extends beyond the consideration of isolated protein targets directly associated with a disease. Instead, it necessitates the consideration of the subnetwork of proteins interacting with the specific target(s) implicated in the disease; this interconnected group is commonly referred to as the “disease module”.[2]

Traditional network models have been widely used to depict intricate interactions within biomedical systems. Although these models have demonstrated impressive capabilities, they often face challenges in capturing the semantic complexity inherent in the diverse relationships among biomedical entities. In response to this limitation, recent approaches have embraced the use of multi-relational networks, such as knowledge graphs (KG).[12]. Defining a knowledge graph (KG) precisely is challenging due to the presence of multiple conflicting definitions in the literature.[13, 14] Here, a specific definition that considers KGs as heterogeneous directed and labeled multigraphs will be adopted, building on top of the one given by the Schlichtkrull et al.[15] Thus, a knowledge graph could be described as a graph-based data structure that contains diverse types of vertices and edges, which can be defined as *G* = (𝒩, ℰ, ℛ, *Ψ*). Each edge within the graph is characterized by a relation type (*r* ∈ ℛ), and represented as a triplet value (*u, r, v*) ∈ ℰ. The vertices, often referred to as entities, are categorized into subsets based on their type (from the set *Ψ*).[14]

KGs serve as structured representations of real-world information. Their ability to model complex, structured data in a machine-readable manner has led to their extensive use in various domains, including question answering, information retrieval, and content-based recommendation systems.[16] Although the initial entity in the triple is commonly known as the head entity, linked to the tail entity through a relation, it’s worth noting that, due to their significant role in encoding human reasoning and language, the components of triplets are often termed as subject, predicate, and object.[15]

There are a lot of different heterogeneous biomedical pharmacological databases representable by KGs, each specializing in a specific domain. These diverse datasets serve a crucial role in progressing biomedical research, education, and diagnostic advancements. Researchers leverage these resources for a multitude of applications, including drug repurposing, as exemplified by Hetionet[17], ParmaKG[12], PharMeBINet[18], and PrimeKG[19]. Despite these endeavors, it’s widely recognized that even the most sophisticated KGs are not complete or perfect.[20, 21] Consequently, researchers have investigated various techniques to fix inaccuracies and fill in missing information within KGs, a task often referred to as Knowledge Graph Completion or Knowledge Graph Augmentation. KG expansion can involve extracting new facts from external sources, experimentally generating new facts, or inferring missing facts based on the existing ones within the KG.[16]

This latter approach, known as Link Prediction (LP), has become an increasingly active research domain, particularly benefiting from the surge in machine learning and deep learning techniques. The majority of LP models leverages KG components to learn low-dimensional representations, commonly referred to as Knowledge Graph Embeddings, then using them to infer new facts.[16]

The impact of deep learning was revolutionary in computer vision [22] and natural language processing [23], yet its applicability remained bounded by the prerequisites of data structure regularity. The convergence of network analysis and machine learning gives rise to graph machine learning (GML), enabling the utilization of graph structures and other non-uniform datasets (such as point clouds, meshes, manifolds, etc.).[24]

At the core of GML methods lies the fundamental concept of acquiring valid feature representations for nodes, edges, or complete graphs. Notably, graph neural networks (GNNs), specialized deep neural network architectures tailored for graph-structured data, systematically enhance the node features within a graph through iterative information propagation from neighboring nodes.[24]

The majority of machine learning techniques applied to graphs are made of two distinct components: a versatile encoder and a task-specific decoder.[25] The encoder is responsible for embedding the graph’s nodes or the entire graph into a feature space of reduced dimensions. For embedding complete graphs, a common approach involves first embedding individual nodes and then applying a permutation invariant pooling function (such as sum, max, or mean) to generate a representation at the graph level. On the other hand, the decoder computes the output that pertains to the specific task under consideration.[24]

Traditional machine learning algorithms usually work by taking a feature vector as input and learning a mapping from this vector to a predictive output. In contrast, incorporating object’s relationships into its representation, Statistical Relational Learning (SRL) is dedicated to constructing statistical models for relational data such as the structures found in Knowledge Graphs. SRL techniques can be applied to existing KGs to develop LP models that predict new facts (triples) by exploiting the underlying information within the existing facts.[26]

Utilizing the previously described notation, where each KG fact is structured as a triple (*h, r, t*) with *h* representing the head (subject), *r* as the relation (predicate), and *t* denoting the tail (object), LP techniques predict the accurate entity to complete (*h, r*, ?) (tail prediction) or (?, *r, t*) (head prediction).[16]

The majority of LP models rely on embeddings to establish a scoring function *f* (***h, r, t***)^*^ for assessing the credibility of a given fact (*h, r, t*). During the prediction phase, when presented with an incomplete triple (*h, r*, ?), the missing tail entity is inferred as the one that, upon inclusion in the triple, produces the highest score:[16]

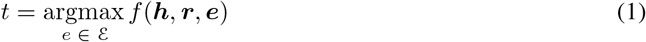

KGEMs generally necessitate the parameterization of all nodes *n* ∈ *N* (entities) and edge types *r* ∈ *R* (relations).[26] Assuming vector embeddings, shallow encoders map these sets to *d*-dimensional vectors *f*_*n*_ : *n* → ℝ^*d*^ and *f*_*r*_ : *r* → ℝ^*d*^. Importantly, these encoders scale linearly with respect to the number of entities and relations, resulting in an entity embedding matrix with *O*(|;*N* |;) space complexity.^*^

This strategy can be effective when applied to small, standard benchmark datasets such as Freebase[27] with approximately 15,000 nodes and WordNet[28] with around 40,000 nodes. However, training on more extensive graphs, such as YAGO 3-10[29] featuring 120,000 nodes or WikiKG2[30] with approximately 2.5 million nodes, presents significant computational challenges.

In a parallel with Natural Language Processing (NLP), shallow node encoding in KGs resembles shallow word embedding, which learns a vocabulary of the most frequent words while treating rarer ones as out-of-vocabulary (OOV). In NLP, the OOV issue has been addressed by enabling the creation of infinite combinations with a finite vocabulary, thanks to subword units. Inspired by this, similar strategies were explored for tokenizing entities within large knowledge graphs (*G* = (𝒩, ℰ, ℛ) constituted of |;𝒩|; nodes, | ℰ |; edges, and |;ℛ|; relation types), resulting in a substantial reduction in parameter complexity, improved generalization, and the natural representation of new, previously unseen entities using a fixed vocabulary. To achieve this, tokenization relies on atomic units analogous to subword units, rather than encompassing the entire set of nodes.

In pursuit of this goal, NodePiece, introduced by Galkin et al.[31], presents an anchor-based method for learning a fixed-size vocabulary 𝒱 (|;𝒱|; ≪ |;𝒩|;) applicable to any connected multi-relational graph. A selected subset of nodes (called anchors, *a* ∈ 𝒜, 𝒜 ⊂ 𝒩) along with all relation types, constitute the set of atoms, which enables the representation of all possible nodes, with the construction of a combinatorial array of sequences from a limited atom vocabulary (𝒱 = 𝒜 ∪ ℛ). In contrast to shallow methods, each node *n* undergoes tokenization^†^, resulting in a unique *hash*(*n*)^‡^ formed by *k* closest anchors, *z*_*a*_^§^ discrete anchor distances and *m* immediate relations (Figure 2). A crucial component for constructing a node embedding is the encoder function *enc*(*n*) : *hash*(*n*) → ℝ^*d*^, which converts the result of the tokenization of a node to its embedding, projecting it from ℝ(*k*+*m*)×^*d*^ to ℝ^*d*^.

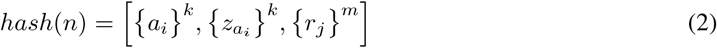

**Figure 1.**
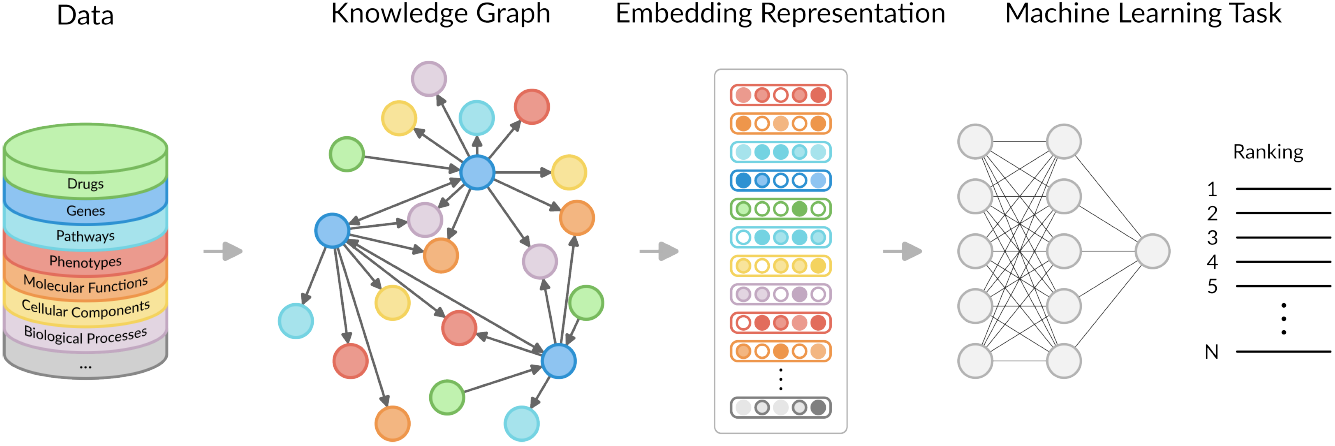
Knowledge Graph Machine Learning. The knowledge graph is constructed using the available data sources. Subsequently, vector representations (embeddings) are learned for entities and relations, which can be employed for a variety of machine learning tasks.

**Figure 2.**
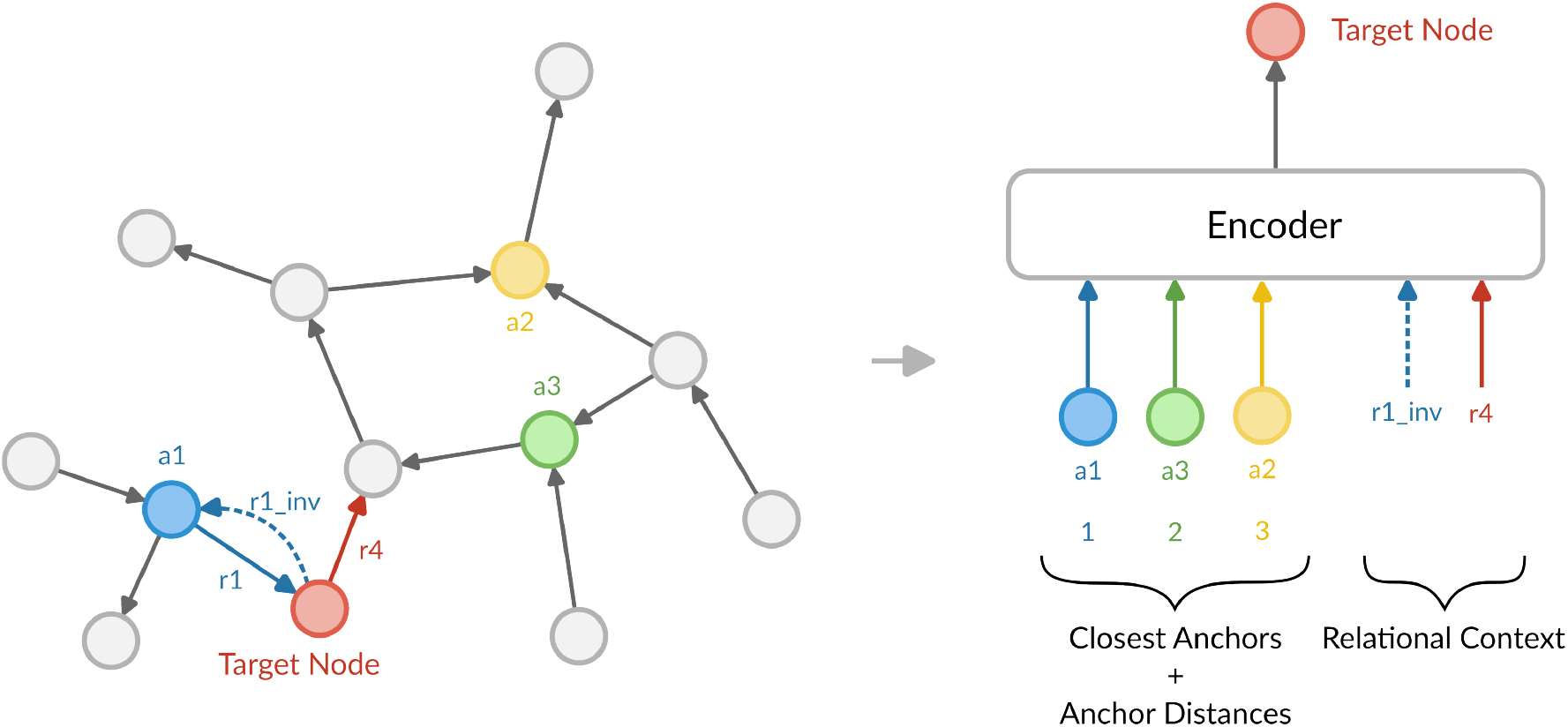
NodePiece Tokenization. Using three anchor points, a1, a2, and a3, a target node is tokenized, resulting in a hash that includes the top-k nearest anchors, their respective distances to the target node, and the relational context of outgoing relations from the target node. This sequence of hashed information is encoded, generating a distinctive embedding. The addition of inverse relations ensures network connectivity. Adapted from Galkin et al.[31]

Taking advantage of this method, the overall parameter allocation is now reduced to a small fixed-size atom vocabulary and the encoder function’s complexity ((*k* + *m*) × *d* instead of |;𝒩|; × *d* + |;ℛ|; × *d*).

Galkin et al.[31] demonstrated that employing a fixed-size NodePiece vocabulary combined with a simple encoder still leads to competitive outcomes across various tasks, encompassing link prediction, node classification, and relation prediction. Additionally, the use of anchor-based hashing enables to operate effectively in both inductive settings and out-of-sample scenarios, accommodating unseen entities during the inference phase.

In this work, we created a comprehensive KG that we named PATHOS (PATHologies of HOmo Sapiens) collecting and integrating data on relevant biological entities from 24 distinct databases. Moreover, we developed LOGOS (Learning Optimized Graph-based representations of Object Semantics), a KGEM capable of providing predictions based on PATHOS.

To evaluate the capabilities of LOGOS, we carried out three crucial case studies: drug repurposing for Alzheimer’s disease (AD), phenotype selection for Huntington’s disease (HD), and the identification of proteins related to multiple sclerosis (MS). These studies showcase the potential of our integrated KG and KGEM to address pressing issues in the field of drug research.

Neurological disorders, like AD, HD, and MS, have profound implications for human health and well-being, resulting in significant suffering and disability. Yet, despite decades of research, effective treatments for these conditions remain elusive. Therefore, our work not only contributes to advancing scientific understanding but also holds the promise of assisting in the development of innovative solutions for these and other diseases.

## 2 Methods

### 2.1 KG Construction (PATHOS)

The analysis of biological systems, diseases, and therapies necessitates the integration of data from various sources. This integration poses unique challenges, principally consisting by the diverse formats and conflicting identifiers employed by each data source. Our approach to addressing these challenges is detailed in the subsequent sections on Data Collection and Data Integration.

#### Data Collection

PATHOS draws upon a collection of data from 24 public databases (see Table 1 and Table S1) renowned for their high-quality structured information pertaining to relevant biological entities.^*^ Our data collection exclusively focused on Homo sapiens.

**Table 1.**
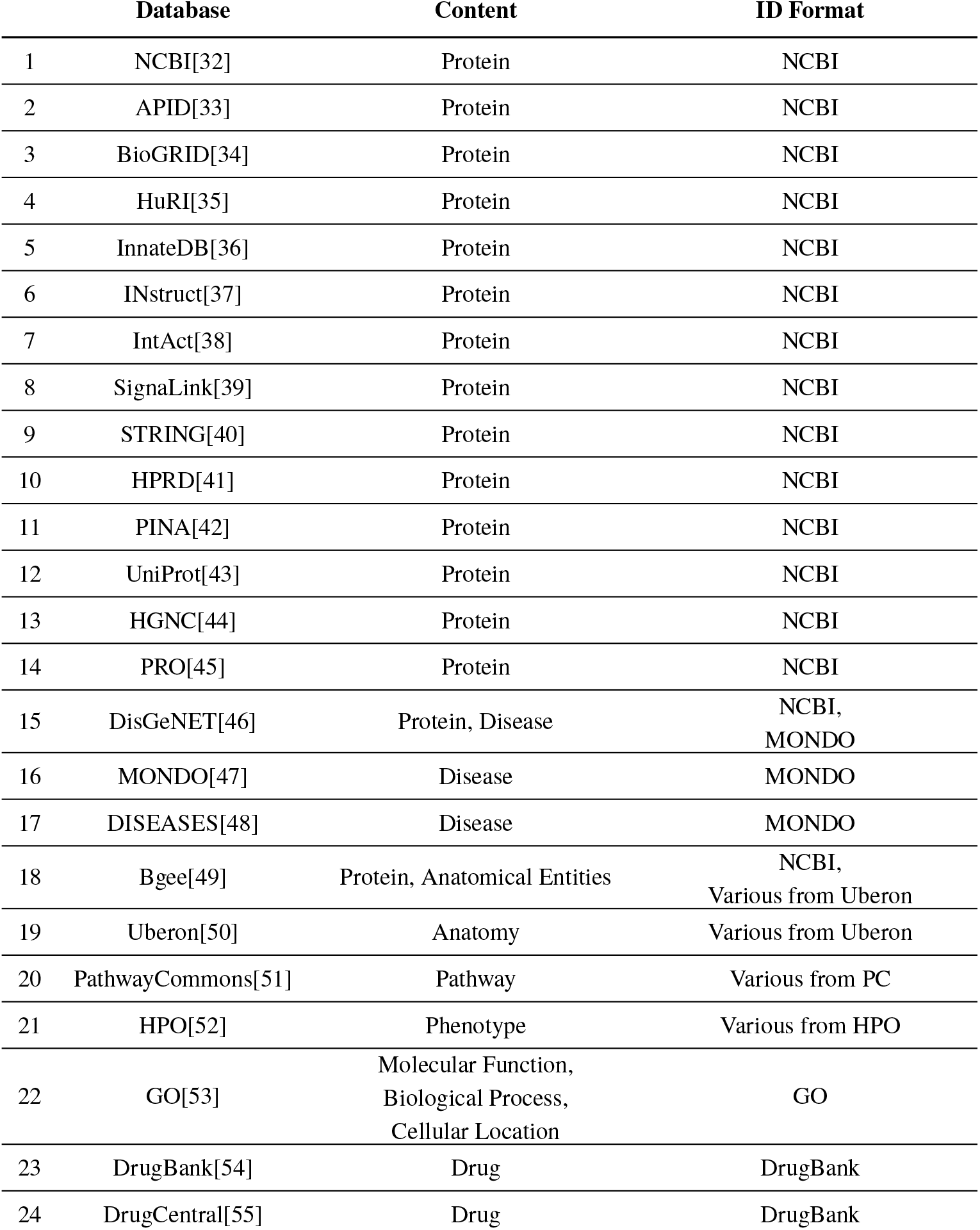
PATHOS Sources. List of all the databases employed for the construction of the KG PATHOS. The main type of content and the relative ID format are also reported. For a list of all the individual files see Table S1.

The source data, stored in a variety of formats, required the development of unique parsers for each data source to enable their transformation into a standardized file format suitable for integration.

#### Data Integration

After standardizing the data, we conducted a merging process, during which duplicate entries were eliminated to prevent redundancy, while maintaining data integrity. This consistency was maintained by mapping all listed entities to official identifiers, including NCBI Entrez ID, MONDO, DrugBank, and more (For the complete reference, please see Table 1).

As a result of this integration effort, we generated a TSV (Tab-separated values) file having eight columns (subject, relation, object, subjectName, objectName, subjectType, objectType, source) and 4,487,349 rows.

### 2.2 KG Embedding Model (LOGOS)

We utilized PyKEEN (Python KnowlEdge EmbeddiNgs, version 1.10.1), an open-source Python package designed for KGEs.[56] This tool enables the construction of KGEMs by offering a diverse range of interaction models, training methodologies, loss functions, and the capacity to explicitly model inverse relations. Indeed, within PyKEEN, a KGEM is constructed as a composite structure with four key components:

#### Interaction Model

we chose NodePiece for its reduced memory footprint and its capability of handling out-of-sample scenarios, as explained above.[31]

#### Loss Function

we applied the self-adversarial negative sampling (NSSA) loss function as proposed by Sun et al.[57], since negative sampling has proven quite effective for KGE[58].

#### Training Approach

in training under the open world assumption (OWA), triples that are not included in the KG are treated as unknown, resulting in over-generalization and poor model performance.[26] Instead, we opted for the stochastic local closed world assumption (sLCWA), which considers a random subset of head and tail generation strategies from LCWA as negative triples. This choice offers several advantages, including reduced computational load, minimal updates to embeddings, and flexibility for integrating new negative sampling strategies.

#### Inverse Relations

we included inverse triples because they are necessary for the employed version of NodePiece to maintain reachability of each node and balance in- and out-degrees.^*^

This extensive toolset within PyKEEN facilitated the development and fine-tuning of LOGOS, our KGEM, to meet the specific needs of our research.

### 2.3 NodePiece

To optimize the hyperparameters for the NodePiece model, we utilized the dedicated function offered by PyKEEN.

Among the most crucial parameters, we employed a 2-layer MLP aggregation encoder, consistent with the NodePiece original paper (ReLU activation and dropout rate of 0.1). The embedding dimension was set at 128, and we worked with 20 entity tokens and 5 relation tokens for each node. Our approach included a total of 10,000 anchors, 80% of them were the top degree nodes and 20% were randomly selected. Negative triples were generated corrupting positive triples with the Bernoulli method[59]. The training loop followed a stochastic local closed-world assumption, and the learning rate was set at 0.0001.

Further details and additional hyperparameters can be found in Supplementary Information.

#### Interaction Functions

Numerous embedding models specific for knowledge graphs have been developed, primarily differing in the scoring of the plausibility of a given triple. In this section, we provide a brief overview of the interaction (scoring) functions explored in our study. These selected functions are well-established in the literature, encompass diverse methodologies, and have started to be investigated within the field of drug discovery.[60]

##### TransE

TransE utilizes a straightforward vector summation in the latent space, where the head entity embedding is added to the relation embedding, bringing the result close to the tail embedding:[61]

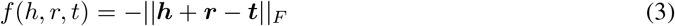

Where *F* is typically either the L1 or L2 norm.^*^

However, this method doesn’t effectively capture one-to-many, many-to-one, and asymmetric relations in practical settings, as the embedding is accurate only when each entity and relation appears in just one fact.

##### DistMult

DistMult employs a vector for each relation type, represented as a diagonal square matrix to significantly reduce the parameter count.[62] However, this means that it is constrained to model symmetric relations exclusively. Its scoring function is:

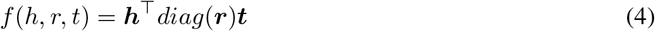

##### ComplEx

In ComplEx, the entity and relation embeddings are complex valued (***h, r, t*** ∈ ℂ^*K*^, K being the dimensionality of the embeddings).[58] Its scoring function becomes:

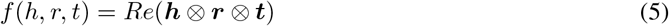

Where *Re*() takes only the real value from the complex number, and ⊗ is the standard componentwise multilinear dot product.^†^[63]

This allows ComplEx to handle both symmetric and asymmetric relations effectively.

##### RotatE

Integrating concepts from various existing models, RotatE employs complex valued embeddings for entities and relations. RotatE forces the modulus of the relation vector to be 1 (∀*i* |;*r*_*i*_|; = 1), and operates in a way that the relation rotates the head to tail entities.[57]

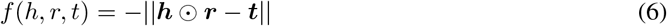

Where ⊙ is the Hadamard product.

This design enables RotatE to handle various relation types, including symmetric, asymmetric, inversion, and composition relations.

### 2.4 Learning Process

The learning process of LOGOS, as a supervised model, begins with the generation of embeddings using a Multilayer Perceptron (MLP). This MLP encodes a hash of tokenized sequences of anchors for each node in the knowledge graph (see Section 1). Subsequently, the model scores both positive and negative triples within the training dataset, with the aim of ranking true triples higher than false triples. Parameters are updated iteratively to minimize the loss during this learning process, and its performance is evaluated on the validation set.

### 2.5 Training Strategy

To ensure robust model performance assessment, we employed a random split of the initial dataset of triples into three distinct subsets: a training set (80% of the data), a validation set (10% of the data), and a test set (10% of the data). This stratified partitioning allowed us to perform rigorous hyperparameter optimization using the training and validation sets. In the end, the model’s performance was evaluated on the test set, which it had never encountered during training.

To account for stochasticity[64], we retrained each of the models five times and evaluated their performance on the same test set. The model that on average exhibited the best performance was employed for the subsequent case studies (complete hyperparameters list is available in Supplementary Information).

### 2.6 Transductive Link Prediction

The task is that of link prediction, which involves the completion of triples of interest either inserting the missing head or tail. Since no new entities are introduced in the triples (the set of nodes in the knowledge graph remains unchanged), this task is considered transductive.

During this phase, LOGOS leverages on the embeddings learned during the training phase to score the triples completed with all the possibile entities in the KG, constructing a ranking based on the results.

### 2.7 Evaluation Protocol

In our evaluation process, we implemented a ranking procedure. To assess the model’s performance, each validation or test triple underwent a corruption process, in which the head entity *h*_*i*_ was removed and replaced by each entity *e*_*i*_ ∈ ℰ\h_i_ in turn. Subsequently, we calculated a score for each triple, sorted these scores in ascending order to determine the rank of the correct triple (*h, r, t*), and then repeated this process for the tail entity as well. The overall model performance was evaluated based on the results of both head and tail corruption procedures (using the mean).

Our evaluation metrics included the mean reciprocal rank (MRR), its adjusted version (AMRR)[65], and the fraction of correct entities in the top-k rank positions (Hits@k) for various values of k: 1, 3, and 10, as well as the adjusted Hits@10.

Additionally, for downstream tasks, we incorporated another widely recognized metric: the areas under the receiver operating characteristic curve (AUC-ROC).

We followed the approach proposed by Bordes et al.[61] ensuring that all corrupted triples did not belong to the original dataset. Additionally, we considered the realistic ranking, for which the rank of an entity is calculated as the mean of the optimistic rank (where the entity is ranked first among those with equal scores) and the pessimistic rank (where the entity is ranked last among those with equal scores), following the method proposed by Berrendorf et al.[65]

### 2.8 Case Studies

In order to demonstrate the capabilities and versatility of PATHOS and LOGOS, we applied them to three distinct case studies in the field of neurological diseases: inferring novel drug candidates for repurposing, selecting plausible phenotypes, and identifying disease-related proteins.

For each case study, we asked LOGOS to complete a triple, either suggesting the subject or the object. In response, it generated a ranked list of entities from the knowledge graph, positioning the most promising candidates at the top. We then verified that the correct entity types were prioritized, assessed the results against the known triples available in PATHOS, and conducted a thorough literature search to find supporting evidence for the top-ranked entities.

#### Drug Repurposing for Alzheimer Disease

We asked LOGOS to complete the triple *(?, indication, Alzheimer’s disease)*. Subsequently, we conducted a literature search for evidence supporting the top 15 suggested drugs.

#### Huntington’s Disease Phenotype Prediction

For this case study, the triple to complete was: *(Huntington disease, has phenotype, ?)*. From the top 50 selected phenotypes, we filtered out those present in the training or validation set, and ensured their consistency with known symptoms and Huntington’s disease phenotypes.

#### Proteins Related to Multiple Sclerosis

In this case study, we aimed to identify the subject of the triple *(?, related to disease, multiple sclerosis)*. To validate the correctness of the answer, we assessed the molecular functions and biological processes enriched in the first 100 proteins within the ranking.

For gene ontology enrichment analysis, we leveraged PANTHER (v 17.0)[66], a powerful and up-to-date tool that seamlessly integrates with the Gene Ontology (GO) website. This system is well-maintained with current GO annotations, ensuring reliable and comprehensive functional annotations. To assure robust results, only gene sets with false discovery rate (FDR) p-values below 0.05 were included in the analysis.

## 3 Results and Discussion

In this section, we will delve into the outcomes of our study, showcasing the twofold contributions of our work: the KG and the KGEM. First, we will provide a descriptive analysis of the knowledge graph, then we will present three compelling case studies, demonstrating the power of our model, focusing on neurological diseases.

Furthermore, we will present the results of our comparative analysis of interaction functions, providing a clear rationale for our selection of the best-performing one for the case studies.

### 3.1 Data Analysis

PATHOS, our comprehensive biomedical knowledge graph, represents a vast network of interconnected information, encompassing 174,367 entities categorized into 17 distinct types. These entities are linked by 4,487,349 relations, spanning 158 unique types (for a complete list check Table S2). Among the most prominent node types within PATHOS are proteins, biological processes, diseases, anatomical entities, molecular functions, phenotypes, and drugs (for a complete list check Table S3). Particularly noteworthy are proteins, diseases and drugs, with 61,862 protein entities, 23,232 disease entities and 8,282 drug entities.

This multifaceted knowledge graph serves as the foundation for our data-driven investigations, enabling us to explore and extract valuable insights in the fields of drug repurposing and disease characterization.

Figure 3 illustrates the metagraph, which is a graphical representation of the relationships and connections within PATHOS. Notably, proteins, as the central entity type with the most nodes and connections, play a pivotal role in shaping the structure of this knowledge graph.

**Figure 3.**
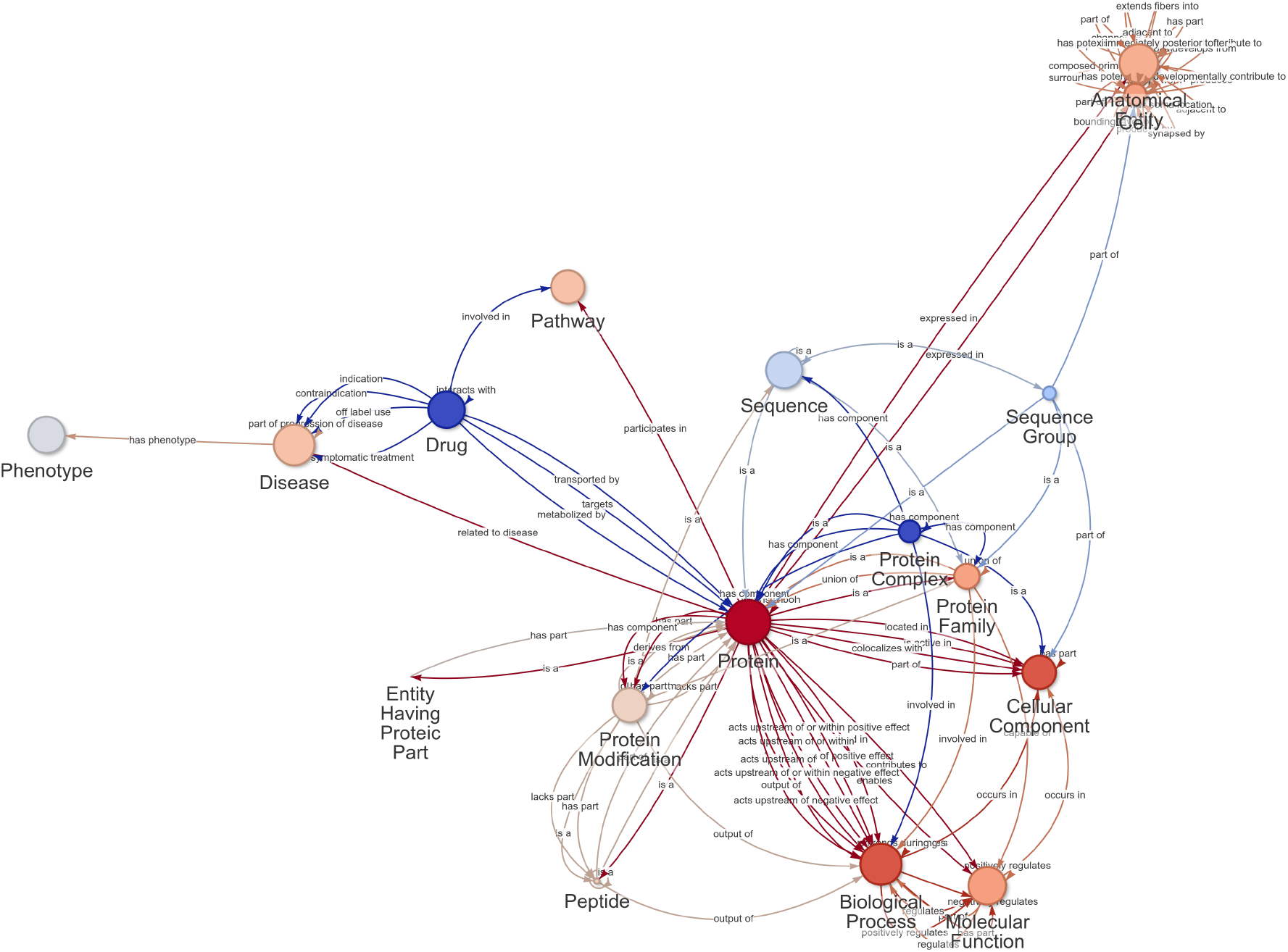
PATHOS Metagraph. This graphical representation illustrates the types of nodes and their possible connections within the PATHOS knowledge graph. Each node represents a type of entity (e.g., proteins, diseases, drugs), and each edge denotes a possible relationship between these entity types. The size of each node is proportional to the number of entities of that type present in the knowledge graph, reflecting their relative abundance.

### 3.2 Comparison of Interaction Functions

Table 2 provides an overview of the performances of several interaction functions, all the models were trained with the same hyperparameters.

**Table 2.**
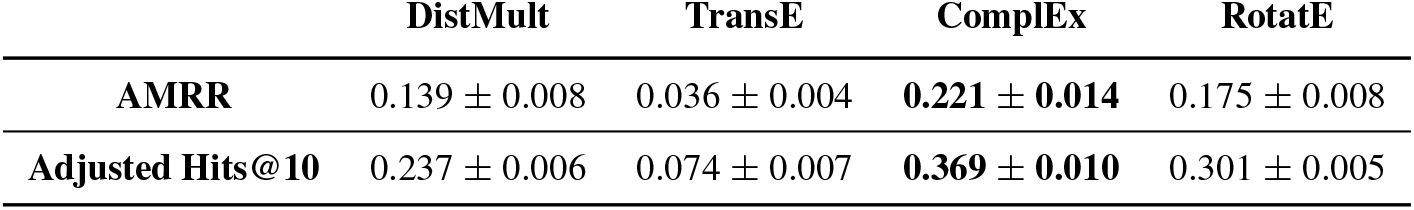
Interaction Functions Comparison. Adjusted mean reciprocal rank and adjusted hits@10 scores for evaluating LOGOS with different interaction functions on the test set. Mean ± standard deviation of 5 replicas.

Surprisingly, the best-performing model, ComplEx, is not the most recent one, RotatE (which is second in line). In light of these results, we selected the ComplEx model with the best AMRR for our subsequent case study predictions.

### 3.3 Drug Repurposing for Alzheimer’s Disease

Our evaluation of LOGOS’s performance involved multiple steps. Initially, we verified its ability to prioritize drugs, which, in this case, is the correct entity type for completing the triple. Remarkably, out of the first 1,000 proposed entities, 997 were indeed drugs, demonstrating LOGOS’s strong capability to prioritize them and achieving a ROC-AUC of 0.99.

Subsequently, we focused on specific predictions for Alzheimer’s disease (AD) indications. For drugs already linked to AD in PATHOS (Epicriptine, Donepezil, Tacrine, Aducanumab, Galantamine, Ipidacrine, Rivastigmine, Acetylcarnitine), the ROC-AUC reached 0.94, considering only the drugs in the ranking (otherwise the ROC-AUC would have been of 1).

Table 3 shows the top 15 highest-scoring novel drug repurposing candidates for AD, including the canonical name of the drug, evidence category and PMID (literature reference supporting the interpretation). Among these top 15 predicted drug candidates, 6 drugs are validated for treating AD based on literature evidence, while 2 candidates exhibit a potential relationship with AD.

**Table 3.**
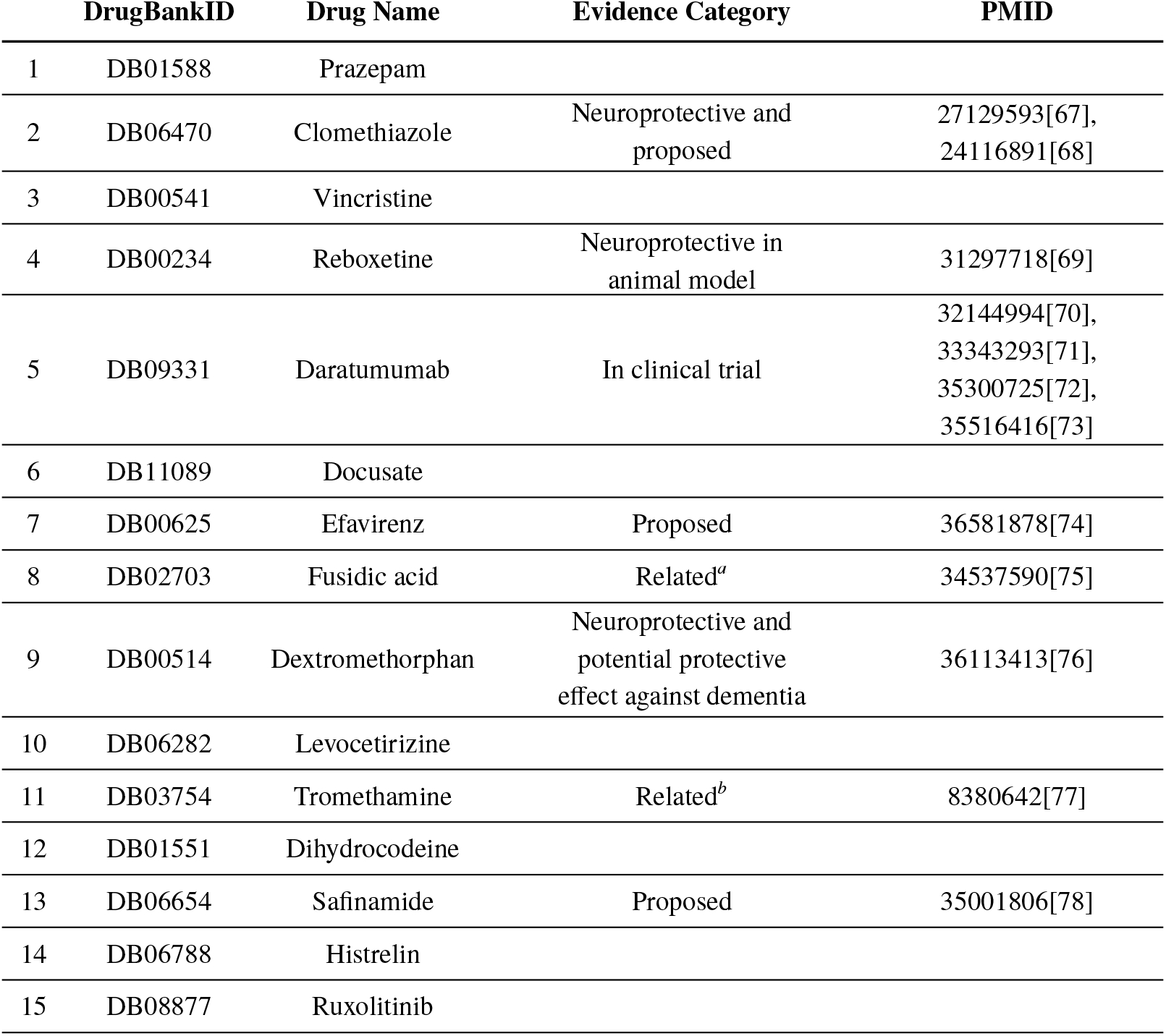
Prioritized Drugs for AD. 15 highest-scoring drug repurposing candidates for Alzheimer’s Disease. It reports: DrugBank ID, canonical name, evidence category and literature reverence. ^a^ Aggregation inhibitor and disaggregator of silk fibroin, it was suggested as a treatment for protein aggregation disorders. ^b^ Blocks amyloid beta channels, a mechanism that was proposed as a useful strategy for drug discovery for treatment of AD

Some noteworthy examples include Daratumumab, Clomethiazole, and Fusidic acid. Daratumumab, an FDA-approved human antibody targeting CD38 for multiple myeloma, is currently in a phase two clinical trial (NCT04070378) for mild to moderate AD due to its immunomodulatory effects on non-plasma cells and its potential to cross the blood-brain barrier.[72] Clomethiazole, an anticonvulsant with demonstrated neuroprotective properties, is considered a promising candidate for future combination therapies addressing neuronal injury[67] and could serve as a lead compound for anti-neurodegenerative drug discovery[68]. Fusidic acid is of particular interest as it aligns with one of the key theories about Alzheimer’s disease etiology: protein aggregation. Indeed, this compound has been suggested as a therapeutic approach for protein aggregation disorders.[75]

### 3.4 Huntington’s Disease Phenotype Prediction

The top 5,967 entities (over 174,367 possible entities) in the ranking were phenotypes, reaffirming LOGOS’s ability to determine the appropriate entity type for completing the triple.

Focusing solely on phenotypes, the ROC-AUC for those already cataloged in PATHOS reached 0.97. Among the first 50 phenotypes (see Table S4), 14 were part of training sets (with none in the validation nor test sets). Impressively, an additional 16 out of the tpo 50 phenotypes matched descriptions found in relevant literature references: dementia, tremor, muscle spasm, abnormality of extrapyramidal motor function, dysarthria, shuffling gait, ataxia, postural tremor, generalized-onset seizure, drooling[79, 80], abnormal autonomic nervous system physiology[81], abnormality of somatosensory evoked potentials[82, 83], muscle weakness[84]. This outcome not only validates LOGOS’s capabilities, but can also suggest potential research avenues for clinical applications, aiming to enhance the anamnesis process.

### 3.5 Proteins Related to Multiple Sclerosis

Out of the entire pool of 174,367 entities, the model correctly prioritized proteins, with the top 17,479 entities in the ranking belonging to this category, underscoring LOGOS efficiency in identifying the appropriate entity type for the task.

For validating the gene ranking, we assessed the ROC-AUC for known ones in PATHOS, resulting in a high score of 0.97.

To provide a more detailed assessment, we examined the first 100 proteins (see list in Table S5) in the ranking, carrying out a gene ontology (GO) enrichment analysis. This analysis revealed strong associations with biological processes (see Table S6) related to MS: chronic inflammation regulation or response (chronic inflammatory response to antigenic stimulus, regulation of chronic inflammatory response to antigenic stimulus), toll-like receptors (toll-like receptor TLR6:TLR2 signaling pathway), macrophage (positive regulation of cellular response to macrophage colony-stimulating factor stimulus), vitamin D (positive regulation of vitamin D biosynthetic process, positive regulation of calcidiol 1-monooxygenase activity). Concerning molecular functions related to MS (see Table S7), the analysis identified relationships with: anandamide (anandamide 11,12 epoxidase activity, anandamide 8,9 epoxidase activity, anandamide 14,15 epoxidase activity)[85, 86], death receptor (death receptor agonist activity)[87], NAD (NAD+ nucleotidase, cyclic ADP-ribose generating, NAD(P)+ nucleosidase activity)[88, 89], MHC class II (MHC class II protein complex binding)[90].

## 4 Limitations

The project, while achieving its core objectives, is not without limitations. Some difficulties are common to many data science initiatives and relate primarily to data availability and model development. For instance, information regarding all biological entities remains unavailable, and the data (and consequently the relations) we do possess is notably skewed towards the most extensively studied entities. Additionally, incoherence of entity identifiers among source databases can result in errors during data integration and linkage. Moreover, the stability and maintenance of the source databases used for PATHOS are essential for keeping it up-to-date. Any disruptions or inconsistencies in these source databases may affect the quality of the KG. While PATHOS is comprehensive, there may still be valuable data types, such as chemical and physical information, side effects or druggability, that are not included. The absence of certain entity and relation types may limit the range of insights and predictions that can be made.

On the model development side, LOGOS was thoughtfully constructed and optimized. However, like any model, there is room for improvement. Potential enhancements include exploring better encoding techniques, increasing embedding size, and optimizing the selection of anchor entities for even more accurate predictions.

In summary, while the project has achieved its primary objectives, it is important to recognize these limitations, and addressing them in the future could further improve the capabilities of PATHOS and LOGOS.

## 5 Conclusions

In conclusion, we gathered and integrated data on relevant biological entities from 24 distinct databases, creating the comprehensive knowledge graph PATHOS.

Additionally, we developed LOGOS, a knowledge graph embedding model, and evaluated its capabilities across three critical tasks: drug repurposing for Alzheimer’s disease, phenotype selection for Huntington’s disease, and the identification of proteins related to multiple sclerosis.

LOGOS succeeded in each of these tasks, demonstrating its potential to help drug research and foster innovative advancements in the field.

## Data and Code Availability

The whole generated data is publicly available from the GitHub repository https://github.com/LucaMenestrina/PATHOS_LOGOS, as well as the full code for the collection, building and analysis. A detailed reference of the source data is provided in the file “data/sources/sources.json” of the aforementioned repository (for every database are reported: name, version, license, employed files, URL and date of access). The Supporting Information can be downloaded at https://github.com/LucaMenestrina/PATHOS_LOGOS/blob/main/SI.pdf.

## Footnotes

* Where ***h, r***, and ***t*** are the embedding vectors of *h, r*, and *t*.

* The emphasis is typically placed on entities since the number of relations |;*R*|; is usually orders of magnitude smaller than |;*E*|;

† Representation of an element with a sequence of other entities called tokens. In NLP, a token is a sequence of characters in some particular document that are grouped together as a useful semantic unit for processing.

‡ A *hash* function is a mathematical function that takes data of variable sizes as input and produces a fixed-size output.

§ Given a graph *G* and a node *n*, the anchor distance 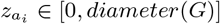 with the anchor node *ai* is defined as the shortest path distance between *a*_*i*_ and *n* in the graph *G*.

* In this paper, we use “protein” to refer specifically to the functional biomolecules that carry out cellular processes, while “gene” denotes the genetic sequences encoding these proteins, underscoring the distinction between the genetic information and its expressed product.

* For each relation *r* ∈ ℛa corresponding inverse relation *r*_*inv*_ is introduced. Consequently, the task of predicting the head entity of a (*r, t*)-pair (thus: (?, *r, t*)) becomes the task of predicting the tail entity of the corresponding inverse pair (*t, r*_*inv*_) (that means: (*t, r*_*inv*_, ?)).

* A norm (|;|;*x*|;|;), in mathematics, is a function mapping elements from a vector (*x*) to non-negative real numbers. In a certain way, it behaves like a distance from the origin. In general, a *p*-norm (L*p*, with *p* a real number *p* ≥ 1) is: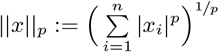 Which means that a L1 norm (the taxicab or Manhattan distance) is: distance) is: 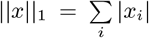 and the L2 (Euclidean distance) is: 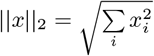

† 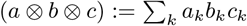

