## Supplementary Information for "Knowledge Graph and Machine Learning Help the Research of Drugs Aimed at Neurological Diseases"

**Luca Menestrina***
Department of Pharmacy and Biotechnology
Alma Mater Studiorum - University of Bologna
Via Belmeloro 6, Bologna 40126, Italy †

**Maurizio Recanatini**
Department of Pharmacy and Biotechnology
Alma Mater Studiorum - University of Bologna
Via Belmeloro 6, Bologna 40126, Italy

### Supporting Information

* Corresponding author.
† Current address: Chemotargets, S.L. Barcelona Science Park (PCB) Baldiri Reixac 4, Barcelona 08028, Catalonia, Spain

**Table S1.** Source Files for PATHOS.

| | Source | License | File | Version |
|---|---|---|---|---|
| 1 | NCBI | Public Domain | `Homo_sapiens.gene_info.gz` | 2023-07-04 (accessed: 2023-07-04) |
| 2 | APID | CC-BY-NC | `9606_Q1.txt` | accessed: 2023-07-04 |
| 3 | BioGRID | MIT | `BIOGRID-ORGANISM-4.4.223.tab.zip` | 4.4.223 (accessed: 2023-07-04) |
| 4 | HuRI | CC BY 4.0 | `HuRI.tsv` | 2020-03-09 (accessed: 2023-07-04) |
| 5 | InnateDB | DESIGN SCIENCE LICENSE | `all.mitab.gz` | 2022-01-29 (accessed: 2023-07-04) |
| 6 | INstruct | All rights reserved (Authorization obtained by e-mail contact with Haiyuan Yu ¡¿) | `sapiens.sin` | 2020-08-13 (accessed: 2021-10-05) |
| 7 | IntAct | CC-BY 4.0 | `intact.zip` | 2023-06-03 (accessed: 2023-07-04) |
| 8 | SignaLink | CC BY-NC-SA 3.0 | `slk3db_dump_json.tgz` | 2022-03-11 (accessed: 2023-07-04) |
| 9 | STRING | CC BY 4.0 | `human.name_2_string.tsv.gz` | 2019-01-27 (accessed: 2023-07-04) |
| 10 | STRING | CC BY 4.0 | `9606.protein.links.full.v11.5.txt.gz` | 2021-10-30 (accessed: 2023-07-04) |
| 11 | HPRD | Freely Available for non-commercial purposes | `HPRD_FLAT_FILES_041310.tar.gz` | 2016-08-20 (accessed: 2022-05-30) |
| 12 | PINA | Freely Downloable All Rights Reserved (check with the develop team https://omics.bjcancer.org/pin a2012/contact.do) | `Homo sapiens-20140521.tsv` | 2014-10-27 (accessed: 2022-05-30) |
| 13 | DisGeNET | CC BY-NC-SA 4.0 | `disease_mappings.tsv.gz` | 2020-05-15 (accessed: 2023-07-04) |
| 14 | DisGeNET | CC BY-NC-SA 4.0 | `curated_gene_disease_associations.tsv.gz` | 2020-05-07 (accessed: 2023-07-04) |
| 15 | MONDO | CC BY 4.0 | `mondo.obo` | 2023-07-03 (accessed: 2023-07-04) |
| 16 | HPO | Freely Available (with conditions) | `hp.obo` | accessed: 2023-07-04 |
| 17 | HPO | Freely Available (with conditions) | `phenotype.hpoa` | accessed: 2023-07-04 |
| 18 | HPO | Freely Available (with conditions) | `genes_to_phenotype.txt` | accessed: 2023-07-04 |
| 19 | DISEASES | CC BY 4.0 | `human_disease_knowledge_filtered.tsv` | 2023-07-02 (accessed: 2023-07-04) |
| 20 | UniProt | CC BY 4.0 | `HUMAN_9606_idmapping.dat.gz` | 2023-06-28 (accessed: 2023-07-04) |
| 21 | PathwayCommons | Freely Available, under the license terms of each contributing database (www.pathwaycommons.org /pc2/datasources) | `PathwayCommons12.All.uniprot.gmt.gz` | 2019-09-18 (accessed: 2023-07-04) |
| 22 | HGNC | Freely Available | `gene_with_protein_product.txt` | 2023-07-03 (accessed: 2023-07-04) |
| 23 | GO | CC BY 4.0 | `goa_human.gaf.gz` | accessed: 2023-07-04 |
| 24 | GO | CC BY 4.0 | `go.obo` | accessed: 2023-07-04 |

| | | | | |
|---|---|---|---|---|
| 25 | PRO | CC BY 4.0 | `pro_reasoned.obo` | 68.0<br>(accessed: 2023-07-04) |
| 26 | Uberon | CC-BY 3.0 | `human-view.obo` | accessed: 2023-07-04 |
| 27 | Bgee | CC0 1.0 | `Homo_sapiens_expr_simple.tsv.gz` | 2021-02-15<br>(accessed: 2021-10-05) |
| 28 | DrugBank | CC BY-NC 4.0 | `all-full-database` | 5.1.10<br>(accessed: 2023-05-23) |
| 29 | DrugCentral | CC BY-SA 4.0 | `drug2disease.tsv` | 2023-05-10<br>(accessed: 2023-07-04) |

**Table S2.** Relation Types in PATHOS.

| Type | # Relations |
|---|---|
| expressed_in | 1965903 |
| interacts_with | 1758237 |
| is_a | 164146 |
| has_phenotype | 128128 |
| participates_in | 81833 |
| involved_in | 80420 |
| related_to_disease | 71506 |
| enables | 52265 |
| located_in | 43361 |
| part_of | 24803 |
| contraindication | 20523 |
| has_symbol | 18803 |
| is_active_in | 16884 |
| targets | 14666 |
| indication | 6798 |
| metabolized_by | 5594 |
| regulates | 3396 |
| negatively_regulates | 2951 |
| positively_regulates | 2945 |
| excluded_subClassOf | 2692 |
| acts_upstream_of_or_within | 1931 |
| off-label use | 1739 |
| lacks_part | 1669 |
| has_part | 1572 |
| develops_from | 1503 |
| has_characteristic | 1292 |
| transported_by | 913 |
| contributes_to | 861 |
| mutually_spatially_disjoint_with | 792 |
| output_of | 773 |
| colocalizes_with | 717 |
| contributes_to_morphology_of | 632 |
| connects | 542 |
| adjacent_to | 350 |
| predisposes_towards | 350 |
| overlaps | 312 |
| disease_has_feature | 279 |
| acts_upstream_of | 257 |
| composed_primarily_of | 240 |
| has_muscle_insertion | 231 |
| connected_to | 225 |
| has_muscle_origin | 219 |
| branching_part_of | 203 |
| has_potential_to_develop_into | 199 |
| has_component | 197 |
| occurs_in | 196 |
| attaches_to | 168 |
| innervated_by | 143 |
| supplies | 140 |
| in_lateral_side_of | 131 |
| extends_fibers_into | 129 |
| continuous_with | 117 |
| innervates | 109 |
| has_developmental_contribution_from | 102 |
| tributary_of | 99 |
| in_left_side_of | 89 |
| drains | 89 |
| luminal_space_of | 87 |
| surrounds | 87 |
| in_right_side_of | 84 |

| | |
|---|---|
| has_skeleton | 79 |
| disease_arises_from_feature | 79 |
| immediate_transformation_of | 71 |
| skeleton_of | 70 |
| derived_from_ancestral_fusion_of | 63 |
| derives_from | 56 |
| proximally_connected_to | 56 |
| conduit_for | 55 |
| bounding_layer_of | 55 |
| disease_has_major_feature | 54 |
| produced_by | 51 |
| disease_shares_features_of | 50 |
| acts_upstream_of_positive_effect | 50 |
| has_muscle_antagonist | 47 |
| transformation_of | 47 |
| immediately_deep_to | 40 |
| channel_for | 36 |
| surrounded_by | 35 |
| existence_ends_during | 34 |
| distally_connected_to | 32 |
| channels_from | 26 |
| capable_of | 25 |
| intersects_midsagittal_plane_of | 25 |
| produces | 25 |
| union_of | 23 |
| has_soma_location | 22 |
| anteriorly_connected_to | 22 |
| developmentally_induced_by | 20 |
| acts_upstream_of_negative_effect | 19 |
| subdivision_of | 19 |
| channels_into | 18 |
| preceded_by | 18 |
| has_member | 17 |
| acts_upstream_of_or_within_positive_effect | 16 |
| superficial_to | 14 |
| existence_starts_during | 14 |
| existence_starts_and_ends_during | 14 |
| dorsal_to | 14 |
| in_posterior_side_of | 14 |
| has_potential_to_developmentally_contribute_to | 13 |
| in_anterior_side_of | 13 |
| immediately_superficial_to | 13 |
| happens_during | 13 |
| ventral_to | 12 |
| anterior_to | 12 |
| sexually_homologous_to | 12 |
| deep_to | 12 |
| preaxialmost_part_of | 11 |
| location_of | 10 |
| contains | 10 |
| develops_in | 9 |
| protects | 9 |
| posterior_to | 9 |
| disease_causes_feature | 8 |
| developmentally_replaces | 8 |
| proximalmost_part_of | 8 |
| immediately_preceded_by | 7 |
| in_superficial_part_of | 7 |
| existence_starts_with | 6 |
| in_deep_part_of | 6 |
| in_dorsal_side_of | 6 |
| postaxialmost_part_of | 5 |
| develops_from_part_of | 5 |
| existence_ends_with | 5 |

| | |
|---|---|
| distalmost_part_of | 5 |
| in_ventral_side_of | 5 |
| filtered_through | 4 |
| indirectly_supplies | 4 |
| anastomoses_with | 4 |
| proximal_to | 4 |
| serially_homologous_to | 3 |
| acts_upstream_of_or_within_negative_effect | 3 |
| precedes | 3 |
| starts | 3 |
| in_innermost_side_of | 2 |
| has_no_connections_with | 2 |
| symptomatic treatment | 2 |
| in_distal_side_of | 2 |
| lumen_of | 2 |
| part_of_progression_of_disease | 2 |
| attaches_to_part_of | 2 |
| in_outermost_side_of | 2 |
| distal_to | 2 |
| in_proximal_side_of | 2 |
| ends | 2 |
| immediately_posterior_to | 1 |
| immediately_anterior_to | 1 |
| ends_with | 1 |
| existence_ends_during_or_before | 1 |
| ends_during | 1 |
| synapsed_by | 1 |
| in_central_side_of | 1 |
| posteriorly_connected_to | 1 |
| existence_starts_during_or_after | 1 |
| directly_develops_from | 1 |
| trunk_part_of | 1 |
| layer_part_of | 1 |
| has_boundary | 1 |

**Table S3.** Node Types in PATHOS.

| Type | # Nodes |
|---|---|
| protein | 58908 |
| biologicalProcess | 27668 |
| disease | 23314 |
| anatomicalEntity | 14288 |
| molecularFunction | 11228 |
| phenotype | 8641 |
| drug | 8282 |
| sequence | 8067 |
| proteinModification | 4954 |
| cellularComponent | 4054 |
| pathway | 3968 |
| proteinFamily | 518 |
| cell | 226 |
| proteinComplex | 221 |
| sequenceGroup | 25 |
| peptide | 4 |
| entityHavingProteicPart | 1 |

#### LOGOS Hyperparameters

**Batch Size** : 256

**Num Epochs** : 100

**Training Loop** : sLCWA

**Optimizer** : Adam

- **Learning Rate** : 0.0001

**Loss** : NSSA

- **Adversarial Temperature** : 0.6868102318671975
- **Margin** : 50

**Model** : NodePiece

- **Aggregation** : MLP
- **Embedding Dimension** : 128
- **Entity Initializer** : Xavier Uniform
- **Interaction** : ComplEx
- **Number of Tokens** : 20, 5
- **Tokenizers** :
  - **Searcher** : ScipySparse
  - **Max Iter** : 100
  - **Selection** : MixtureAnchorSelection
  - **Number of Anchors** : 10,000
  - **Ratios** : 0.8, 0.2
  - **Selections** : Degree, Random

**Negative Sampler** : Bernoulli

- **Number of Negatives per Positive** : 100

**Table S4.** First 50 Phenotypes Selected for Huntington's Disease.

| | ID | Name | Train Set | Val Set | Test Set |
|---|---|---|---|---|---|
| 1 | HP:0030015 | Female anorgasmia | ✗ | ✗ | ✗ |
| 2 | HP:0002072 | Chorea | ✓ | ✗ | ✗ |
| 3 | HP:0003324 | Generalized muscle weakness | ✓ | ✗ | ✗ |
| 4 | HP:0000716 | Depression | ✓ | ✗ | ✗ |
| 5 | HP:0000741 | Apathy | ✓ | ✗ | ✗ |
| 6 | HP:0002307 | Drooling | ✗ | ✗ | ✗ |
| 7 | HP:0002340 | Caudate atrophy | ✓ | ✗ | ✗ |
| 8 | HP:0002362 | Shuffling gait | ✗ | ✗ | ✗ |
| 9 | HP:0002174 | Postural tremor | ✗ | ✗ | ✗ |
| 10 | HP:0002529 | Neuronal loss in central nervous system | ✓ | ✗ | ✗ |
| 11 | HP:0002460 | Distal muscle weakness | ✗ | ✗ | ✗ |
| 12 | HP:0002071 | Abnormality of extrapyramidal motor function | ✗ | ✗ | ✗ |
| 13 | HP:0001283 | Bulbar palsy | ✗ | ✗ | ✗ |
| 14 | HP:0012332 | Abnormal autonomic nervous system physiology | ✗ | ✗ | ✗ |
| 15 | HP:0001336 | Myoclonus | ✓ | ✗ | ✗ |
| 16 | HP:0002151 | Increased serum lactate | ✗ | ✗ | ✗ |
| 17 | HP:0001332 | Dystonia | ✓ | ✗ | ✗ |
| 18 | HP:0002921 | Abnormal cerebrospinal fluid morphology | ✗ | ✗ | ✗ |
| 19 | HP:0001288 | Gait disturbance | ✓ | ✗ | ✗ |
| 20 | HP:0008652 | Autonomic erectile dysfunction | ✗ | ✗ | ✗ |
| 21 | HP:0001260 | Dysarthria | ✗ | ✗ | ✗ |
| 22 | HP:0003387 | Decreased number of large peripheral myelinated nerve fibers | ✗ | ✗ | ✗ |
| 23 | HP:0006801 | Hyperactive deep tendon reflexes | ✗ | ✗ | ✗ |
| 24 | HP:0000726 | Dementia | ✗ | ✗ | ✗ |
| 25 | HP:0002063 | Rigidity | ✓ | ✗ | ✗ |
| 26 | HP:0002922 | Increased CSF protein concentration | ✗ | ✗ | ✗ |
| 27 | HP:0002197 | Generalized-onset seizure | ✗ | ✗ | ✗ |
| 28 | HP:0030319 | Weakness of facial musculature | ✗ | ✗ | ✗ |
| 29 | HP:0012751 | Abnormal basal ganglia MRI signal intensity | ✗ | ✗ | ✗ |
| 30 | HP:0001251 | Ataxia | ✗ | ✗ | ✗ |
| 31 | HP:0012416 | Hypercapnia | ✗ | ✗ | ✗ |
| 32 | HP:0100021 | Cerebral palsy | ✗ | ✗ | ✗ |
| 33 | HP:0000738 | Hallucinations | ✓ | ✗ | ✗ |
| 34 | HP:0003394 | Muscle spasm | ✗ | ✗ | ✗ |
| 35 | HP:0025331 | Upgaze palsy | ✗ | ✗ | ✗ |
| 36 | HP:0007377 | Abnormality of somatosensory evoked potentials | ✗ | ✗ | ✗ |
| 37 | HP:0012670 | Orthostatic syncope | ✗ | ✗ | ✗ |

| | | | | | |
|---|---|---|---|---|---|
| 38 | HP:0001337 | Tremor | ✗ | ✗ | ✗ |
| 39 | HP:0011289 | EEG with temporal sharp slow waves | ✗ | ✗ | ✗ |
| 40 | HP:0001324 | Muscle weakness | ✗ | ✗ | ✗ |
| 41 | HP:0002120 | Cerebral cortical atrophy | ✗ | ✗ | ✗ |
| 42 | HP:0000737 | Irritability | ✓ | ✗ | ✗ |
| 43 | HP:0009045 | Exercise-induced rhabdomyolysis | ✗ | ✗ | ✗ |
| 44 | HP:0000488 | Retinopathy | ✗ | ✗ | ✗ |
| 45 | HP:0002141 | Gait imbalance | ✓ | ✗ | ✗ |
| 46 | HP:0000739 | Anxiety | ✓ | ✗ | ✗ |
| 47 | HP:0000511 | Vertical supranuclear gaze palsy | ✗ | ✗ | ✗ |
| 48 | HP:0000802 | Impotence | ✗ | ✗ | ✗ |
| 49 | HP:0410263 | Brain imaging abnormality | ✗ | ✗ | ✗ |
| 50 | HP:0040141 | Tardive dyskinesia | ✗ | ✗ | ✗ |

**Table S5.** First 100 Proteins Related to Multiple Sclerosis.

| | ID | Name | Train Set | Val Set | Test Set |
|---|---|---|---|---|---|
| 1 | TTR | transthyretin | ✗ | ✗ | ✗ |
| 2 | ALB | albumin | ✗ | ✗ | ✗ |
| 3 | TSNAX-DISC1 | TSNAX-DISC1 readthrough (NMD candidate) | ✗ | ✗ | ✗ |
| 4 | MIR885 | microRNA 885 | ✗ | ✗ | ✗ |
| 5 | POMC | proopiomelanocortin | ✓ | ✗ | ✗ |
| 6 | SHBG | sex hormone binding globulin | ✗ | ✗ | ✗ |
| 7 | ADIPOQ | adiponectin, C1Q and collagen domain containing | ✗ | ✗ | ✗ |
| 8 | SLC10A2 | solute carrier family 10 member 2 | ✗ | ✗ | ✗ |
| 9 | CYP2D6 | cytochrome P450 family 2 subfamily D member 6 | ✗ | ✗ | ✗ |
| 10 | TF | transferrin | ✗ | ✗ | ✗ |
| 11 | MIR99A | microRNA 99a | ✗ | ✗ | ✗ |
| 12 | MIR346 | microRNA 346 | ✗ | ✗ | ✗ |
| 13 | CNR2 | cannabinoid receptor 2 | ✗ | ✗ | ✗ |
| 14 | MIR505 | microRNA 505 | ✗ | ✗ | ✗ |
| 15 | CYP2C8 | cytochrome P450 family 2 subfamily C member 8 | ✗ | ✗ | ✗ |
| 16 | TNF | tumor necrosis factor | ✗ | ✗ | ✗ |
| 17 | CP | ceruloplasmin | ✗ | ✗ | ✗ |
| 18 | CYP2B6 | cytochrome P450 family 2 subfamily B member 6 | ✗ | ✗ | ✗ |
| 19 | VEGFA | vascular endothelial growth factor A | ✗ | ✗ | ✗ |
| 20 | ACE2 | angiotensin converting enzyme 2 | ✗ | ✗ | ✗ |
| 21 | GSTM1 | glutathione S-transferase mu 1 | ✗ | ✗ | ✗ |
| 22 | IL6 | interleukin 6 | ✗ | ✗ | ✗ |

| | | | | | |
|---|---|---|---|---|---|
| 23 | RBP4 | retinol binding protein 4 | ✗ | ✗ | ✗ |
| 24 | MIR412 | microRNA 412 | ✗ | ✗ | ✗ |
| 25 | CYP2E1 | cytochrome P450 family 2 subfamily E member 1 | ✗ | ✗ | ✗ |
| 26 | TLR4 | toll like receptor 4 | ✗ | ✗ | ✗ |
| 27 | MIR433 | microRNA 433 | ✗ | ✗ | ✗ |
| 28 | SLC30A6 | solute carrier family 30 member 6 | ✗ | ✗ | ✗ |
| 29 | MIR766 | microRNA 766 | ✗ | ✗ | ✗ |
| 30 | MIR192 | microRNA 192 | ✗ | ✗ | ✗ |
| 31 | VKORC1 | vitamin K epoxide reductase complex subunit 1 | ✗ | ✗ | ✗ |
| 32 | MIR218-1 | microRNA 218-1 | ✗ | ✗ | ✗ |
| 33 | HMOX1 | heme oxygenase 1 | ✗ | ✗ | ✗ |
| 34 | DAOA | D-amino acid oxidase activator | ✗ | ✗ | ✗ |
| 35 | MTHFR | methylenetetrahydrofolate reductase | ✗ | ✗ | ✗ |
| 36 | TYK2 | tyrosine kinase 2 | ✓ | ✗ | ✗ |
| 37 | HLA-DQA2 | major histocompatibility complex, class II, DQ alpha 2 | ✗ | ✗ | ✗ |
| 38 | GJB5 | gap junction protein beta 5 | ✗ | ✗ | ✗ |
| 39 | MIR17 | microRNA 17 | ✗ | ✗ | ✗ |
| 40 | IL10 | interleukin 10 | ✗ | ✗ | ✗ |
| 41 | MIR98 | microRNA 98 | ✗ | ✗ | ✗ |
| 42 | PTGS2 | prostaglandin-endoperoxide synthase 2 | ✗ | ✗ | ✗ |
| 43 | PSCA | prostate stem cell antigen | ✗ | ✗ | ✗ |
| 44 | EDN1 | endothelin 1 | ✗ | ✗ | ✗ |
| 45 | IGHG1 | immunoglobulin heavy constant gamma 1 (G1m marker) | ✗ | ✗ | ✗ |
| 46 | EPHX2 | epoxide hydrolase 2 | ✗ | ✗ | ✗ |
| 47 | FGF2 | fibroblast growth factor 2 | ✗ | ✗ | ✗ |
| 48 | FAXDC2 | fatty acid hydroxylase domain containing 2 | ✗ | ✗ | ✗ |
| 49 | TRH | thyrotropin releasing hormone | ✗ | ✗ | ✗ |
| 50 | OXT | oxytocin/neurophysin I prepropeptide | ✗ | ✗ | ✗ |
| 51 | SCN10A | sodium voltage-gated channel alpha subunit 10 | ✗ | ✗ | ✗ |
| 52 | ERGIC3 | ERGIC and golgi 3 | ✗ | ✗ | ✗ |
| 53 | MIR296 | microRNA 296 | ✗ | ✗ | ✗ |
| 54 | TFF2 | trefoil factor 2 | ✗ | ✗ | ✗ |
| 55 | MIR3622B | microRNA 3622b | ✗ | ✗ | ✗ |
| 56 | INS | insulin | ✗ | ✗ | ✗ |
| 57 | LRP2 | LDL receptor related protein 2 | ✗ | ✗ | ✗ |
| 58 | PLCG2 | phospholipase C gamma 2 | ✗ | ✗ | ✗ |
| 59 | NGF | nerve growth factor | ✗ | ✗ | ✗ |
| 60 | THBD | thrombomodulin | ✗ | ✗ | ✗ |

| | | | | | |
|---|---|---|---|---|---|
| 61 | TLR6 | toll like receptor 6 | ✗ | ✗ | ✗ |
| 62 | MIR30B | microRNA 30b | ✗ | ✗ | ✗ |
| 63 | FCGR1A | Fc gamma receptor Ia | ✗ | ✗ | ✗ |
| 64 | SCD | stearoyl-CoA desaturase | ✗ | ✗ | ✗ |
| 65 | WT1 | WT1 transcription factor | ✗ | ✗ | ✗ |
| 66 | HLA-DPB1 | major histocompatibility complex, class II, DP beta 1 | ✗ | ✗ | ✗ |
| 67 | MPO | myeloperoxidase | ✗ | ✗ | ✗ |
| 68 | GC | GC vitamin D binding protein | ✗ | ✗ | ✗ |
| 69 | SH2B3 | SH2B adaptor protein 3 | ✗ | ✗ | ✗ |
| 70 | IGF2 | insulin like growth factor 2 | ✗ | ✗ | ✗ |
| 71 | PRKCQ | protein kinase C theta | ✗ | ✗ | ✗ |
| 72 | IFNG | interferon gamma | ✗ | ✗ | ✗ |
| 73 | SLC22A1 | solute carrier family 22 member 1 | ✗ | ✗ | ✗ |
| 74 | PROC | protein C, inactivator of coagulation factors Va and VIIIa | ✗ | ✗ | ✗ |
| 75 | UGT1A1 | UDP glucuronosyltransferase family 1 member A1 | ✗ | ✗ | ✗ |
| 76 | FXYD6 | FXYD domain containing ion transport regulator 6 | ✗ | ✗ | ✗ |
| 77 | HP | haptoglobin | ✗ | ✗ | ✗ |
| 78 | SERPINA1 | serpin family A member 1 | ✗ | ✗ | ✗ |
| 79 | HLA-DRB1 | major histocompatibility complex, class II, DR beta 1 | ✗ | ✗ | ✓ |
| 80 | MIR218-2 | microRNA 218-2 | ✗ | ✗ | ✗ |
| 81 | TLR2 | toll like receptor 2 | ✗ | ✗ | ✗ |
| 82 | PPARG | peroxisome proliferator activated receptor gamma | ✗ | ✗ | ✗ |
| 83 | ZAP70 | zeta chain of T cell receptor associated protein kinase 70 | ✗ | ✗ | ✗ |
| 84 | UCN | urocortin | ✗ | ✗ | ✗ |
| 85 | CHRNB2 | cholinergic receptor nicotinic beta 2 subunit | ✗ | ✗ | ✗ |
| 86 | MIR708 | microRNA 708 | ✗ | ✗ | ✗ |
| 87 | MSR1 | macrophage scavenger receptor 1 | ✗ | ✗ | ✗ |
| 88 | ATP1B2 | ATPase Na+/K+ transporting subunit beta 2 | ✗ | ✗ | ✗ |
| 89 | ABCB1 | ATP binding cassette subfamily B member 1 | ✗ | ✗ | ✗ |
| 90 | APOC3 | apolipoprotein C3 | ✗ | ✗ | ✗ |
| 91 | SLCO1B1 | solute carrier organic anion transporter family member 1B1 | ✗ | ✗ | ✗ |
| 92 | HELLPAR | HELLP associated long non-coding RNA | ✗ | ✗ | ✗ |
| 93 | MIR629 | microRNA 629 | ✗ | ✗ | ✗ |
| 94 | LRP1 | LDL receptor related protein 1 | ✗ | ✗ | ✗ |
| 95 | MS4A1 | membrane spanning 4-domains A1 | ✗ | ✗ | ✗ |
| 96 | CCR2 | C-C motif chemokine receptor 2 | ✗ | ✗ | ✗ |

| | | | | | |
|---|---|---|---|---|---|
| 97 | C2 | complement C2 | X | X | X |
| 98 | IFNB1 | interferon beta 1 | X | X | X |
| 99 | EPO | erythropoietin | X | X | X |
| 100 | RNASE3 | ribonuclease A family member 3 | X | X | X |

**Table S6.** First 10 Enriched Biological Processes.

| | ID | Label | Fold Enrichment | FDR |
|---|---|---|---|---|
| 1 | GO:0002439 | chronic inflammatory response to antigenic stimulus | 242.26 | 0.0028 |
| 2 | GO:0038124 | toll-like receptor TLR6:TLR2 signaling pathway | 242.26 | 0.0028 |
| 3 | GO:1990268 | response to gold nanoparticle | 242.26 | 0.0028 |
| 4 | GO:1903974 | positive regulation of cellular response to macrophage colony-stimulating factor stimulus | 242.26 | 0.0028 |
| 5 | GO:0042496 | detection of diacyl bacterial lipopeptide | 242.26 | 0.0027 |
| 6 | GO:0060557 | positive regulation of vitamin D biosynthetic process | 242.26 | 0.0027 |
| 7 | GO:0017187 | peptidyl-glutamic acid carboxylation | 161.51 | 0.0042 |
| 8 | GO:1904466 | positive regulation of matrix metallopeptidase secretion | 161.51 | 0.0042 |
| 9 | GO:0002874 | regulation of chronic inflammatory response to antigenic stimulus | 161.51 | 0.0042 |
| 10 | GO:0060559 | positive regulation of calcidiol 1-monooxygenase activity | 161.51 | 0.0042 |

**Table S7.** First 10 Enriched Molecular Functions.

| | ID | Label | Fold Enrichment | FDR |
|---|---|---|---|---|
| 1 | GO:0062188 | anandamide 11,12 epoxidase activity | 161.51 | 0.0239 |
| 2 | GO:0062187 | anandamide 8,9 epoxidase activity | 161.51 | 0.0233 |
| 3 | GO:0038177 | death receptor agonist activity | 121.12 | 0.0321 |
| 4 | GO:0062189 | anandamide 14,15 epoxidase activity | 121.12 | 0.0313 |
| 5 | GO:0061809 | NAD+ nucleotidase, cyclic ADP-ribose generating | 45.42 | 0.0112 |
| 6 | GO:0050135 | NAD(P)+ nucleosidase activity | 45.42 | 0.0104 |
| 7 | GO:0008392 | arachidonic acid epoxygenase activity | 45.42 | 0.0101 |
| 8 | GO:0023026 | MHC class II protein complex binding | 35.89 | 0.0022 |
| 9 | GO:0070330 | aromatase activity | 27.95 | 0.0310 |
| 10 | GO:0005179 | hormone activity | 18.93 | 0.0000 |
